# Responses to heartbeats in ventromedial prefrontal cortex contribute to subjective preference-based decisions

**DOI:** 10.1101/776047

**Authors:** Damiano Azzalini, Anne Buot, Stefano Palminteri, Catherine Tallon-Baudry

## Abstract

Forrest Gump or Matrix? Preference-based decisions are subjective and entail self-reflection. However, these self-related features are unaccounted for by known neural mechanisms of valuation and choice. Self-related processes have been linked to a basic interoceptive biological mechanism, the neural monitoring of heartbeats, in particular in ventromedial prefrontal cortex (vmPFC), a region also involved in value encoding. We thus hypothesized a functional coupling between the neural monitoring of heartbeats and the precision of value encoding in vmPFC. Human participants were presented with pairs of movie titles. They indicated either which movie they preferred, or performed a control objective visual discrimination that did not require self-reflection. Using magnetoencephalography, we measured heartbeat-evoked responses (HERs) before option presentation, and confirmed that HERs in vmPFC were larger when preparing to the subjective, self-related task. We retrieved the expected cortical value network during choice with time-resolved statistical modeling. Crucially, we show that larger HERs before option presentation are followed by stronger value encoding during choice in vmPFC. This effect is independent of the overall vmPFC baseline activity. The neural interaction between HERs and value encoding predicted preference-based choice consistency over time, accounting for both inter-individual differences and trial-to-trial fluctuations within individuals. Neither cardiac activity nor arousal fluctuations could account for any of the effects. HERs did not interact with the encoding of perceptual evidence in the discrimination task. Our results show that the self-reflection underlying preference-based decisions involves the integration of HERs to subjective value encoding in vmPFC, and that this integration contributes to preference stability.

**Significance statement:** Deciding whether you prefer Forrest Gump or Matrix is based on subjective values, which only you, the decision-maker, can estimate and compare, by asking yourself. Yet, how self-reflection is biologically implemented and its contribution to subjective valuation are not known. We show that in ventromedial prefrontal cortex, the neural response to heartbeats, an interoceptive self-related process, influences the cortical representation of subjective value. The neural interaction between the cortical monitoring of heartbeats and value encoding predicts choice consistency, i.e. whether you consistently prefer Forrest Gump over Matrix over time. Our results pave the way for the quantification of self-related process in decision making and may shed new light on the relationship between maladaptive decisions and impaired interoception.

## Introduction

Do you prefer Forrest Gump or Matrix? The decision is subjective: only *you* know which movie you like best. The subjective values used in preference-based decision making are internally generated, intrinsically private, and entail self-reflection. In other words, estimating a subjective value requires a reflection about how an item affects *you*. In contrast, the evidence required to decide which of the two words ‘listen’ and ‘look’ has more characters is publicly and objectively available to any reader of this article. While the neural underpinnings of valuation and choice have been well studied, the biological mechanism supporting the self-reflection intrinsic to subjective decisions remains unspecified. It might derive from the simplest biological implementation of self-reflection, i.e., the monitoring of one’s current physiological state (Craig, 2002; Blanke and Metzinger, 2009; Damasio, 2010; Park and Tallon-Baudry, 2014; Azzalini et al., 2019) required to select the most appropriate behavior to restore homeostatic balance thus ensuring the integrity of the living organism. It follows that the organism needs to track its internal state to assign a value to a given option (Keramati and Gutkin, 2014; Juechems and Summerfield, 2019), and that an imprecise representation of the internal state may lead to suboptimal choice (Paulus, 2007).

The monitoring of current physiological state is notably indexed by the transient neural response automatically elicited by each heartbeat, also known as the heartbeat-evoked response (HER)(Montoya et al., 1993; Kern et al., 2013). HERs have been linked to subjective, self-related cognitive processes in ventromedial prefrontal cortex (vmPFC)(Park et al., 2014; Babo-Rebelo et al., 2016a, 2016b). A separate stream of studies repeatedly showed that vmPFC encodes subjective values (Lebreton et al., 2009; Bartra et al., 2013; Grueschow et al., 2015). We thus hypothesized that (a) HERs in vmPFC would signal the recruitment of self-reflective processes in preparation to a subjective decision, but absent when preparing to an objective decision and (b) that HER fluctuations would affect valuation in subjective preference-based decisions, but not in decisions based on objective evidence publicly available in the outside world, such as perceptual discriminations. We tested these hypotheses in a paradigm where participants performed either a subjective, preference-based choice, or a control, objective perceptual discrimination, between two visually presented movie titles (**Fig. 1**), while their neural and cardiac activity were measured with magnetoencephalography (MEG) and electrocardiography (ECG), respectively. Each trial began with an instruction period, with a symbol indicating which type of decision to perform, during which we measured HERs. Once options were displayed, participants selected the title of the movie they preferred in subjective preference trials, and the title written with the highest contrast in objective perceptual discrimination trials. We found that (a) HERs during the instruction period were larger when preparing for preference-based decisions than for discrimination ones, and we demonstrated that (b) HER amplitude interacted with the neural encoding of subjective value in vmPFC during choice. The neural interaction between HER and value encoding was associated with more consistent subjective choices. This functional coupling was specific to subjective decisions: HER did not interact with the encoding of perceptual evidence in objective visual discrimination trials.

**Figure 1.**
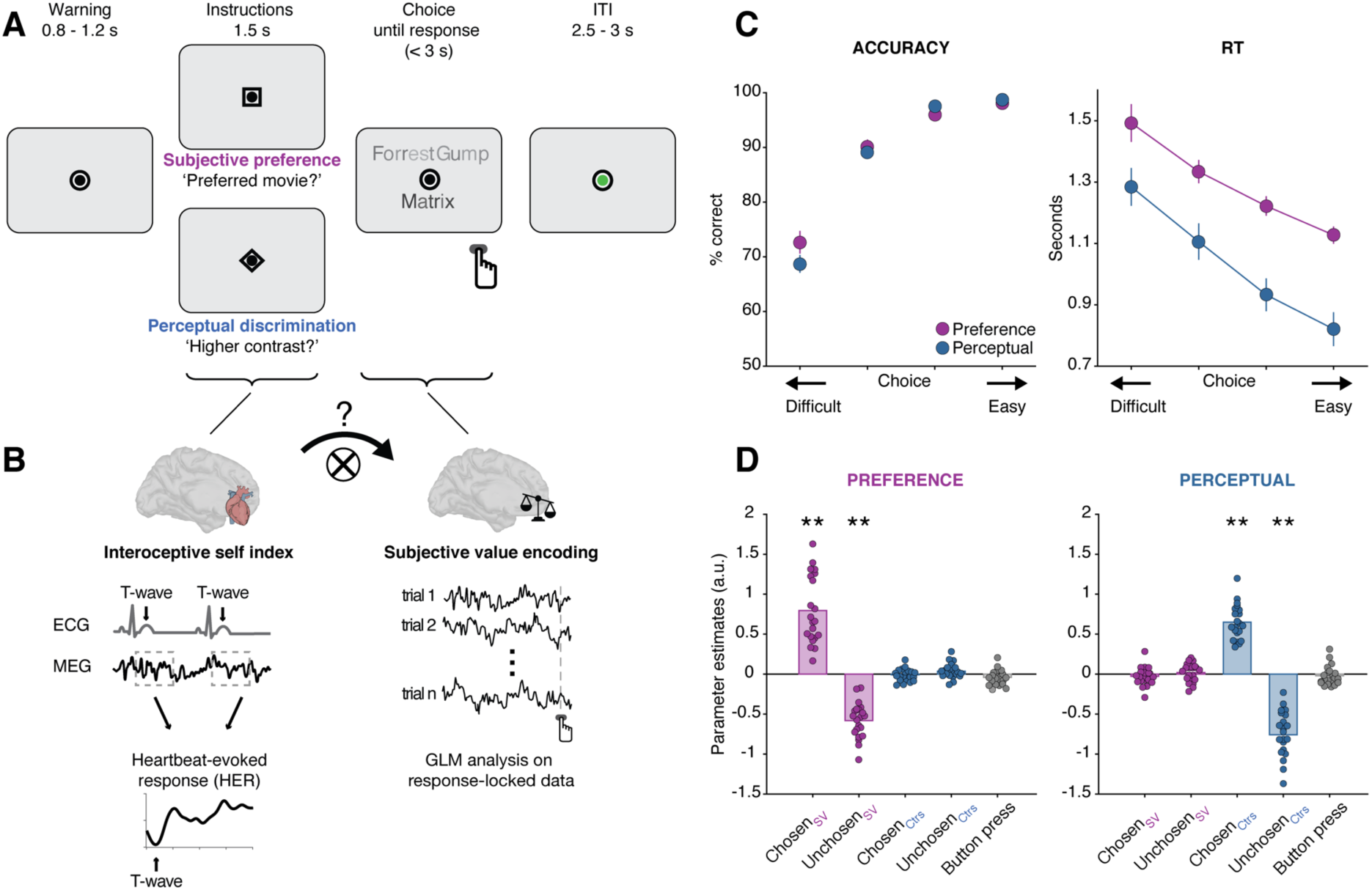
Experimental design and behavioral results. **A**, Trial time-course. After a fixation period of variable duration, a symbol (square or diamond) instructed participants on the type of decision to perform on the upcoming movie titles, either a subjective preference-based choice or an objective perceptual discrimination task. Decision type varied on a trial-by-trial basis. The two movie titles appeared above and below fixation and remained on screen until response or until 3 seconds had elapsed. **B**, Rationale for data analysis. Left, interoceptive self-related processes were indexed by heartbeat-evoked responses (HERs) computed during the instruction period, before option presentation, by averaging MEG activity locked to the T-waves of the electrocardiogram. Right, response-locked MEG activity during choice was modeled on a trial-by-trial basis with a GLM to isolate the spatio-temporal patterns of neural activity encoding value. The central question is whether HERs before option presentation and value encoding interact. **C**, Behavioral results. In both tasks, performance (choice consistency or discrimination accuracy) increased (F_(3,60)_ = 99.25, p < 10^−15^) and response times decreased (F_(3,60)_ = 41.14, p < 10^−13^) for easier choices (i.e. larger difference in subjective value for preference-based decisions, or difference in contrast for perceptual ones). **D**, Only task-relevant information significantly contributed to choice in both preference-based and perceptual decisions as estimated by logistic regression (two-tailed t-test against zero). Each dot represents one participant. **p<0.01.

## Materials and Methods

### Participants

24 right-handed volunteers with normal or corrected-to-normal vision took part in the study after having given written informed consent. They received monetary compensation for their participation. The ethics committee CPP Ile de France III approved all experimental procedures. Three subjects were excluded from further analysis: one subject for too low overall performance (74%, 2 standard deviations below the mean = 87.6%), one subject was excluded for excessive number of artifacts (29.7% of trials, above 2 standard deviations, mean = 7%), one subject was excluded because the ICA correction of the cardiac artifact was not successful.

21 subjects were thus retained for all subsequent analyses (9 male; mean age: 23.57 ± 2.4 years; mean ± SD).

### Tasks and procedure

Participants came on two consecutive days to the lab (mean elapsed time between the two sessions 22.28 ± 3.55 hours) to complete two experimental sessions. The first session was a likability rating on movies (behavior only), from which we drew the stimuli used in the second experimental session, during which brain activity was recorded with magnetoencephalography (MEG).

#### Rating session

We selected 540 popular movies form the Web (allocine.fr) whose title maximal length was 16 characters (spaces included). DVD covers and titles of the pre-selected movies were displayed one by one on a computer screen and subjects had to indicate whether they had previously watched the currently displayed movie by pressing a ‘yes’ or ‘no’ key on a computer keyboard, without any time constraint. Participants were then presented with the list of movie titles they had previously watched and asked to name the 2 movies they liked the most and the two they liked the least. Participants were explicitly instructed to use these 4 movies as reference points (the extremes of the rating scale) to rate all other movies. Last, the titles and the covers of the movies belonging to the list were displayed one by one at the center of the computer screen in random order. Participants assigned to each movie a likability rating by displacing (with arrow keys) a cursor on a 21-points Likert scale and validated their choice with an additional button press. Likability ratings were self-paced and the starting position of the cursor was randomized at every trial.

#### Stimuli

Experimental stimuli consisted of 256 pairs of written movie titles drawn from the list of movies that each participant had rated on the first day. Each movie title was characterized along two experimental dimensions: its likability rating (as provided by the participant) and its contrast. The mean contrast was obtained by averaging the luminance value (between 40 and 100; grey background at 190) randomly assigned to each character of the title. We manipulated trials difficulty by pairing movie titles so that the differences between the two items along the two dimensions (i.e., likability and contrast) were parametric and orthogonal. Additionally, we controlled that the sum of ratings and the sum of contrast within each difficulty level was independent of their difference and evenly distributed. Each pair of stimuli was presented twice in the experiment: one per decision type. A given movie title could appear in up to 10 different pairs. The position of the movie titles on the screen was pseudo-randomly assigned so that the position of the correct option (higher likeability rating or higher contrast) was fully counter-balanced.

#### Experimental task

On the second day, subjects performed a two-alternative forced-choice (2AFC) task while brain activity was recorded with MEG. At each trial, participants were instructed to perform one of the two decisions types on the pair of movie titles (**Fig. 1A**): either a preference decision, in which they had to indicate the item they liked the most, or a perceptual discrimination, in which they had to indicate the title written with the higher contrast. Each trial began with a fixation period of variable duration (uniformly distributed between 0.8 and 1.2 seconds in step of 0.05 s) indicated by a black fixation dot surrounded by a black ring (internal dot, 0.20° of visual angle; external black ring, 0.40° of visual angle), starting from which participants were required not to blink anymore. Next, the outer ring of the fixation turned either into a square or a diamond (0.40° and 0.56° visual angle, respectively) indicating which type of decision participants were to perform (preference-based or perceptual, counter-balanced across participants), for 1.5 seconds. Then, the outer shape turned again into a ring and two movie titles appeared above and below it (visual angle 1.09°). Options remained on screen until response was provided (via button press with the right hand) or until 3 seconds had elapsed. After response delivery, movie titles disappeared and the black fixation dot surrounded by the black circle remained on screen for 1 more second. The central dot turned green and stayed on screen for variable time (uniformly distributed between 2.5 and 3 seconds in step of 0.05 s), indicating participants that they were allowed to blink before the beginning of next trial. Each recording session consisted of 8 blocks of 64 trials each.

Prior to the recording session, participants familiarized themselves with the experimental task by carrying out 3 training blocks. The first 2 blocks (10 trials each) comprised trials of one type only, hence preceded by the same cue symbol. The last block contained interleaved trials (n=20), as in the actual recording. The movie pairs used during training were not presented again during the recording session.

#### Heartbeat counting task

After performing the 8 experimental blocks, we assessed participants’ interoceptive abilities by asking them to count their heartbeats by focusing on their bodily sensations, while fixating the screen (Schandry, 1981). Subjects performed six blocks of different durations (30, 45, 60, 80, 100, 120 seconds) in randomized order. No feedback on performance was provided. Since the acquisition of our dataset, this widely used paradigm has been criticized in several respects (Ring et al., 2015; Desmedt et al., 2018; Zamariola et al., 2018).

#### Questionnaires

Once subjects left the MEG room, they filled 4 questionnaires in French: Beck’s Depression Inventory (BDI) (Beck et al., 1961), Peter’s et al. Delusions Inventory (PDI) (Peters et al., 2004), the Trait Anxiety Inventory (STAI) (Spielberger et al., 1983) and the Obsessive-Compulsive Inventory (OCI) (Foa et al., 2002).

### Recordings

Neural activity was continuously recorded using a MEG system with 102 magnetometers and 204 planar gradiometers (Elekta Neuromag TRIUX, sampling rate 1000 Hz, online low-pass filter at 330 Hz). Cardiac activity was simultaneously recorded (BIOPAC Systems, Inc.; sampling frequency 1000 Hz; online filter 0.05-35 Hz). The electrocardiogram was obtained from 4 electrodes (2 placed in over the left and right clavicles, 2 over left and right supraspinatus muscles (Gray et al., 2007)) and referenced to another electrode on the left iliac region of the abdomen, corresponding to four vertical derivations. The four horizontal derivations were computed offline by subtracting the activity of two adjacent electrodes. Additionally, we measured beat-to-beat changes in cardiac impedance, to compute the beat-by-beat stroke volume (i.e. the volume of blood ejected by the heart at each heartbeat (Kubicek et al., 1970)). Impedance cardiography is a non-invasive technique based on the impedance changes in the thorax due to the changes in fluid volume (blood). A very low-intensity (400 µA rms) high frequency (12.5 kHz) electric current was injected via two source electrodes: the first one was placed on the left side of the neck and the second 30 cm below it (roughly on the sixth rib). Two other monitoring electrodes (placed 4 cm apart from the source ones: below the source electrode on the neck and above the source electrode on the rib cage) measured the voltage across the tissue. To determine left ventricular ejection time, aortic valve activity was recorded by placing an a-magnetic homemade microphone (online band-pass filter 0.05-300 Hz) on the chest of the subject.

Pupil diameter and eye movements were tracked using an eye-tracker device (EyeLink 1000, SR Research) and 4 electrodes (2 electrodes placed on the left and right temples and 2 electrodes placed above and below participant’s dominant eye).

### Cardiac events and parameters

Cardiac events were detected on the right clavicle-left abdomen ECG derivation in all participants. We computed a template of the cardiac cycle, by averaging a subset of cardiac cycles, which was then convolved with the ECG time series. R-peaks were identified as peaks of the result of the convolution, normalized between 0 and 1, exceeding 0.6. All other cardiac waves were defined with respect R-peak. In particular, T-waves were identified as the maximum amplitude occurring within 420 milliseconds after the Q-wave. R-peak and T-wave automatic detection was visually verified for each participant.

Inter-beat intervals (IBIs) were defined for each phase of the trial as the intervals between two consecutive R-peaks. More specifically, we considered for ‘fixation’, ‘instruction period’ and ‘response’ phases the two R-peaks around their occurrence. IBIs during ‘choice’ were based on the two R-peaks preceding response delivery. Inter-beat variability was defined as the standard deviation across trials of IBIs in a given trial phase.

Stroke volume was computed according to the formula (Kubicek et al., 1970; Sherwood et al., 1990):

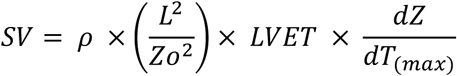

where *ρ* is the resistivity of the blood (135 Ohms*cm) (Berntson et al., 2007), *L*^2^ is the distance between the two source electrodes, *Zo*^2^ is the base impedance, *LVET* is the systolic left ventricular ejection time (in seconds), *dZ*/*dT*_(*max*)_ is the largest impedance change during systole (Ohms/sec). Note that because we obtained stroke volume by injecting a current at 12.5 kHz, rather than the more typical frequency of 100 kHz, absolute stroke volumes are systematically underestimated, but relative values are preserved.

**MEG data preprocessing**

External noise was removed from the continuous data using MaxFilter algorithm. Continuous data were then high-pass filtered at 0.5 Hz (4^th^ order Butterworth filter). Trials (defined as epochs ranging from fixation period to 1 second after response) contaminated by muscle and movement artifacts were manually identified and discarded from further analyses (6% of trials on average, ranging from 0% to 15%).

Independent component analysis (ICA) (Delorme and Makeig, 2004), as implemented in FieldTrip Toolbox (Oostenveld et al., 2011) was used to attenuate the cardiac artifact on MEG data. ICA was computed on MEG data epoched ±200 ms around the R-peak of the ECG, in data segments that were free of artifacts, blinks and saccades above 3 degrees. The number of independent components to compute was set to be equal to the rank of the MEG data. Mean pairwise phase consistency (PPC) was estimated for each independent component (Vinck et al., 2010) with the right clavicle-left abdomen ECG derivation signal in the frequency band 0-25 Hz. Components (up to 3) that exceeded 3 standard deviations from mean PPC were then removed from the continuous data.

To correct for blinks, 2-seconds segments of data were used to estimate blink and eye-movement components. Mean PCC was then computed with respect to vertical EOG signal, and components exceeding mean PCC + 3 standard deviations were removed from continuous data. The procedure was iterated until no component was beyond 3 standard deviations or until 3 components in total were removed. Stereotypical blink components were manually selected in two participants as the automated procedure failed to identified them.

ICA-corrected data were then low-pass filtered at 25 Hz (6^th^ order Butterworth filter)

### Trials selection

Trials had to meet the following criteria to be included in all subsequent analysis: no movement artifacts, sum of blinking periods less than 20% of total trial time, at least one T-peak during instruction period (cf. HERs section), and reaction time neither too short (at least 250 ms) nor too long. To identify exceedingly long RTs, we binned the trials of each task in 4 difficulty levels based on the difference of the two options (i.e., difference in ratings in preference-based choice and difference in contrast for the perceptual ones). Within each difficulty level, for correct and error trials separately, we excluded the trials with reaction times exceeding the participant’s mean RT + 2 standard deviations.

The average number of trials retained per participant was 421.67 ± 43.36 (mean ± SD).

### Heartbeat-evoked responses

Heartbeat-evoked responses were computed on MEG data time-locked to T-wave occurring during the instruction period. T-waves had to be at minimum 400 ms distance from the subsequent R-peak. In order to avoid contamination by transient visual responses or by preparation to the subsequent visual presentation, we only retained T-waves that occurred at least 300 milliseconds after the onset of the instruction cue and 350 milliseconds before the onset of options presentation. If more than one T-wave occurred in this period, HERs for that trial were averaged. HERs were analyzed from T-wave + 50 ms to minimize contamination by the residual cardiac artifact (Dirlich et al., 1997) after ICA correction.

We verify that differences in HERs between the two types of decision were truly locked to heartbeats, and that a difference of similar magnitude could not arise by locking the data to any time point of the instruction period. To this end, we created surrogate timings for heartbeats (within the instruction period), to break the temporal relationship between neural data and heartbeats, and computed surrogate HERs. We created 500 surrogate heartbeat data set, by permuting the timings of the real T-wave between trials belonging to the same decision type (i.e., the timing of the T-wave at trial *i* was randomly assigned to trial *j*). We then searched for surrogate HER differences between trial types using a cluster-based permutation test (Maris and Oostenveld, 2007) (see below). For each of the 500 iterations, we retained the value of the largest cluster statistics (sum(t)) to estimate the distribution of the largest difference that could be obtained randomly sampling ongoing neural activity during the same instruction period. To assess statistical significance, we compared the cluster statistics from the original data against the distribution of surrogate statistics.

### Nonparametric statistical testing of MEG data

HERs difference between preference-based and perceptual trials during instruction presentation was tested for statistical significance using cluster-based permutation two-tailed t-test (Maris and Oostenveld, 2007) as implemented in FieldTrip toolbox (Oostenveld et al., 2011), on magnetometer activity in the time-window 50-300 ms after T-wave. This method defines candidate clusters of neural activity based on spatio-temporal adjacency exceeding a statistical threshold (p < 0.05) for a given number of neighboring sensors (n=3). Each candidate cluster is assigned a cluster-level test statistics corresponding to the sum of *t* values of all samples belonging to the given cluster. The null distribution is obtained non-parametrically by randomly shuffling conditions labels 10,000 times, computing at each iteration the cluster statistics and saving the largest positive and negative *t* sum. Monte Carlo p value corresponds to the proportion of cluster statistics under the null distribution that exceed the original cluster-level test statistics. Because the largest chance values are retained to construct the null distribution, this method intrinsically corrects for multiple comparisons across time and space. Controls analyses involving the clustering procedure were performed with the same parameters.

The significance of beta time-series obtained from GLM analyses at the sensor level was obtained using cluster-based permutation two-tailed t-test against zero.

### Bayes factor

We used Bayes factors (BF) to quantify the evidence in support of the null hypothesis (H_0_ = no difference between 2 measures). To this aim, we computed the maximum log-likelihood of a gaussian model in favor of the alternative hypothesis and for the model favoring the null adjusting the effect size to correspond to a p = 0.05 for our sample size (n = 21 for all analyses except for pupil for which n = 16 and for 3 ECG derivations for which n=20). Finally, we computed Bayesian information criterion and the corresponding Bayes factor. As a summary indication, BF < 0.33 provides substantial evidence in favor of the null hypothesis, BF between 0.33 and 3 does not provide enough evidence for or against the null (Kass and Raftery, 1995).

For regression analyses, Bayes Factor was computed using the online calculator tool (http://pcl.missouri.edu/bf-reg) based on Liang and colleagues (Liang et al., 2008).

### Generalized linear model on response-locked single trials

To analyze how task-related variables are encoded in neural activity during decision, we ran a generalized linear model (GLM) on baseline-corrected (−500 to −200 ms before instruction presentation) single trial MEG data time-locked to button press. We predicted z-scored MEG activity at each time-point and channel using task-relevant experimental variables. For preference-based decisions we modeled MEG activity as:

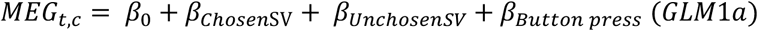

where *t* and *c* represent MEG activity at time-point *t* at channel *c, β*_0_ is the intercept, *β*_*ChosenSV*_ are the z-scored ratings of the chosen option, *β*_*UnchosenSV*_ is the z-scored rating of the alternative unchosen option and *β*_*Button press*_ is a categorical variable representing motor response (i.e. top or bottom).

Similarly, for perceptual decisions we used:

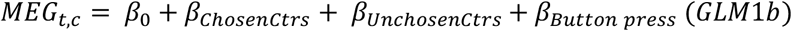

where *β*_*ChosenCtrs*_ and *β*_*UnchosenCtrs*_ are the z-scored contrast of the chosen and unchosen option, respectively.

This procedure provided us with time series of beta values at each channel that could be tested against zero for significance using spatio-temporal clustering (Maris and Oostenveld, 2007). Once significant clusters encoding task-related variables were identified at the sensor level, we reconstructed the cortical sources corresponding to the sensor-level activity averaged within the significant time-window. We modeled source-reconstructed neural activity with the same GLMs to identify the cortical areas mostly contributing to the significant sensor-level effect.

### Generalized linear model on posterior right vmPFC

To quantify the influence of HER in anterior r-vmPFC during instructions on subjective value encoding during choice, we modeled the activity of posterior r-vmPFC, encoding subjective value with the following GLM:

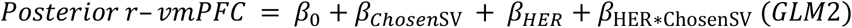

where, *β*_0_ is the intercept, *β*_*ChosenSV*_ are the z-scored ratings of the chosen option, *β*_*HER*_ is the z-scored activity in the anterior right vmPFC cluster defined by comparing HERs in preference-based vs. perceptual choices and *β*_*ChosenSV*HER*_ is the interaction term obtained by multiplying the z-scored previous predictors.

To verify that the interaction between subjective value encoding and HER amplitude was specifically time-locked to heartbeats and not a general influence of baseline activity in anterior r-vmPFC, we ran an alternative model to explain posterior r-vmPFC activity:

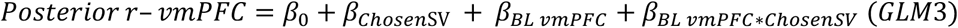

where, *β*_0_ is the intercept, *β*_*ChosenSV*_ are the z-scored ratings of the chosen option, *β*_*BL vmPFC*_ is the z-scored activity in anterior r-vmPFC during instructions averaged across the whole instruction period, not time-locked to heartbeats and *β*_*BL vmPFC*\**ChosenSV*_ is the interaction between the two preceding predictors. Note that regressors are not orthogonalized in any of the GLMs.

### Anatomical MR acquisition and preprocessing

An anatomical T1 scan was acquired for each participant on a 3 Tesla Siemens TRIO (n = 2) or Siemens PRISMA (n = 20) or Siemens VERIO (n = 2). Cortical segmentation was obtained by using automated procedure as implemented in the FreeSurfer software package (Fischl et al., 2004). The results were visually inspected and used for minimum-norm estimation.

### Source reconstruction

Cortical localization of neural activity was performed with BrainStorm toolbox (Tadel et al., 2011). After co-registration of individual anatomy and MEG sensors, 15,003 current dipoles were estimated using a linear inverse solution from time-series of magnetometers and planar gradiometers (weighted minimum-norm, SNR of 3, whitening PCA, depth weighting of 0.5) using overlapping-spheres head model. Current dipoles were constrained to be normally oriented to cortical surface, based on individual anatomy. Source activity was obtained by averaging sensor-level time-series in the time-windows showing significant effects (difference between HERs and beta values different from zero), spatially smoothed (FWHM 6 mm) and projected onto standard brain model (ICBM152_T1, 15,003 vertices). Note that sources in subcortical regions cannot be retrieved with the reconstruction method used here.

To assess which cortical areas contributed the most to the effects observed at the sensor-level, we ran parametric two-tailed t-test and reported all clusters of activity spatially extending more than 20 vertices with individual t-values corresponding to p < 0.005 (uncorrected for multiple comparisons). We reported the coordinates of vertices with the maximal t value and their anatomical labels according to AAL atlas (Tzourio-Mazoyer et al., 2002). For clusters falling into prefrontal cortices, we reported the corresponding areas according to the connectivity-based parcellation developed by Neubert and colleagues (Neubert et al., 2015).

### Pupil data analysis

Pupil data that contained blinks (automatically detected with EyeLink software and extended before and after by 150 ms), saccades beyond 2 degrees and segments in which pupil size changed abruptly (signal temporal derivative exceeding 0.3, arbitrary unit) were linearly interpolated. All interpolated portions of the data that exceeded 1 second were removed from further analyses. Continuous pupil data from each experimental block were then band-pass filtered between 0.01 and 10 Hz (second order Butterworth) and z-scored. 16 subjects were retained for pupil analysis; 5 subjects were excluded due to too low quality of data. Pupil analysis was performed in two ways: 1) averaged pupil diameter in the same time period used for HER computation (i.e., 300 ms after instruction presentation until 350 ms before options display) and 2) averaged pupil diameter in the time-window spanning 1 second before button press until its execution.

### Code accessibility

The custom code and the source data used for the main analyses of this paper can be accessed online at https://github.com/DamianoAzzalini/HER-preferences. Participants did not give any formal agreement to publicly share the MEG and physiological data, hence the raw data supporting the findings are available from the corresponding authors upon reasonable request.

## Results

### Behavioral results

Participants were asked to choose between two simultaneously presented movie titles according either to their subjective preferences or to the visual contrast of movie titles, as indicated by trial-by-trial instructions presented before the alternatives (**Fig. 1A**). Decision difficulty, operationalized as the difference between the two options (in the preference task: difference between likeability ratings measured one day before the MEG session [see Material and Methods], in the perceptual task: difference between contrasts), had the expected impact on behavior in both tasks. Both preference consistency and discrimination accuracy increased and reaction times decreased for easier decisions (**Fig. 1C**; preference task, one-way repeated measure ANOVA, main effect of difficulty: accuracy, F_(3,60)_ = 99.25, p < 10^−15^; RT, F_(3,60)_ = 41.14, p < 10^−13^; perceptual task, main effect of difficulty: accuracy, F_(3,60)_ = 280.2, p < 10^−15^; RT, F_(3,60)_ = 87.67, p < 10^−15^). Preference and perceptual decisions were matched in accuracy (two-way repeated measures ANOVA, main effect of task on accuracy, F_(1,20)_ = 0.38, p = 0.55, interaction task x difficulty, F_(3,60)_ = 2.53, p = 0.07), but preference-based decisions were generally slower (two-way repeated measures ANOVA, main effect of task, RT, F_(1,20)_ = 57.64, p < 10^−6^) and reaction times decreased less rapidly for easier decisions (interaction task x difficulty, F_(3,60)_ = 4.08, p = 0.01). Participants only used task-relevant information (subjective value or objective contrast) to decide, since non-relevant information could not predict choice (**Fig. 1D**; **Table 1**).

**Table1.** Logistic regression against choice, for task-relevant and task-irrelevant stimulus information.

| **Regressor** | **Preference decisions** | **Perceptual decisions** |
| --- | --- | --- |
|  | **two-tailed t-test against 0** | **two-tailed t-test against 0** |
|  | **(uncorrected p)** | **(uncorrected p)** |
| Chosen<sub>SV</sub> | $t_{(20)} = 8.59, p = 3.8 \times 10^{-8}$ | $t_{(20)} = -1.14, p = 0.27$ |
| Unchosen<sub>SV</sub> | $t_{(20)} = -11.95, p = 1.46 \times 10^{-10}$ | $t_{(20)} = 0.78, p = 0.45$ |
| Chosen<sub>Ctr</sub> | $t_{(20)} = -0.50, p = 0.62$ | $t_{(20)} = 13.65, p = 1.36 \times 10^{-11}$ |
| Unchosen<sub>Ctr</sub> | $t_{(20)} = 1.83, p = 0.08$ | $t_{(20)} = -12.17, p = 1.05 \times 10^{-10}$ |
| Button Press | $t_{(20)} = -1.88, p = 0.07$ | $t_{(20)} = -0.79, p = 0.44$ |

### Neural responses to heartbeats are larger when preparing for preference-based decisions

HERs in vmPFC have been previously shown to index self-related processes (Park et al., 2014; Babo-Rebelo et al., 2016a, 2016b). We thus predicted that HERs would be larger when preparing for subjective, preference-based decisions relying on self-reflection than when preparing for objective, perceptual ones. Using a non-parametric clustering procedure which corrects for multiple comparisons across time and space (Maris and Oostenveld, 2007), we found that HERs during the instruction period, before option presentation, were indeed larger when participants prepared for preference-based decisions than for perceptual ones (**Fig. 2A, 2B**; non-parametric clustering, 201-262 ms after T-wave, sum(t) = 1789, Monte Carlo cluster level p = 0.037). Averaging cluster activity separately in the two conditions results in an effect size Cohen’s d of 1.28. The cortical regions that mostly contributed to this effect (**Fig. 2C**) were localized as expected in right and left anterior vmPFC (areas 11m and 14 bilaterally; cluster peak at MNI coordinates, [1 57 −21] and [−3 47 −6]; t-values, 4.68 and 3.76, respectively), but also in the right post-central complex ([32 −22 56]; t-value = 5.57) and right supramarginal gyrus ([41 −33 43]; t-value = 3.98).

**Figure 2.**
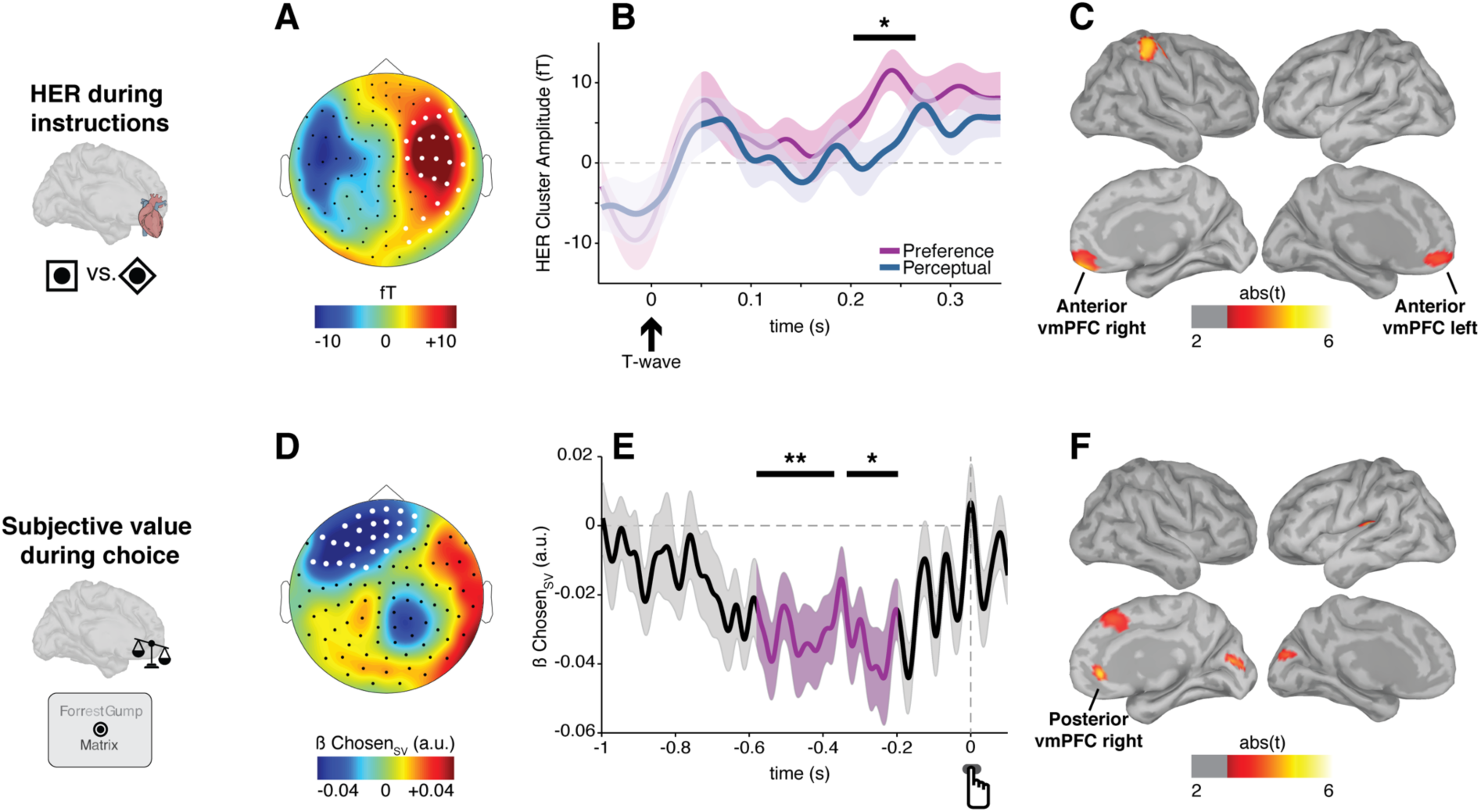
HERs and subjective value encoding. **A**, Topography of the significant HER difference between preference-based and perceptual decisions during the instruction period (201-262 ms after cardiac T-wave, cluster Monte Carlo p = 0.037). **B**, Time-course of HER (± SEM) for preference-based and perceptual decisions in the cluster highlighted in white in A. The portion of the signal (50 ms after T-wave) still potentially contaminated by the cardiac artifact appears in lighter color. The black bar indicates the significant time-window, as established by non-parametric clustering procedure. **C**, Brain regions mostly contributing to the HER difference between preference-based and perceptual decisions (at least 20 contiguous vertices with uncorrected p < 0.005). **D**, Topography of the significant encoding of the chosen option subjective value (−580 to −197 ms before motor response) during choice in preference-based trials. **E**, Time-course (± SEM) of the GLM parameter estimate for the chosen option subjective value in the cluster highlighted in white in D. Black bars indicate significant time-windows, as established by non-parametric clustering procedure. **F**, Brain regions mostly contributing to the encoding of the subjective value of the chosen option (at least 20 contiguous vertices with uncorrected p < 0.005). *p<0.05, **p<0.01.

The HER difference between subjective preference-based trials and objective perceptual discrimination trials was not accompanied by any difference in ECG activity (paired t-test on 4 ECG vertical derivations, all p ≥0.89, all BF≤0.24; paired t-test on 4 ECG horizontal derivations, all p≥ 0.34, all BF ≤ 0.47), in cardiac parameters (inter-beat intervals, inter-beat intervals variability, stroke volume) or arousal indices (alpha power and pupil diameter) measured during the instruction period (**Table 2**). The HER difference is thus neither due to differences in cardiac inputs nor to overall changes in brain state. Importantly, the HER difference was time-locked to heartbeats and thus did not reflect a baseline difference between conditions (Monte-Carlo p=0.026. see **Methods** for details).

**Table 2.**
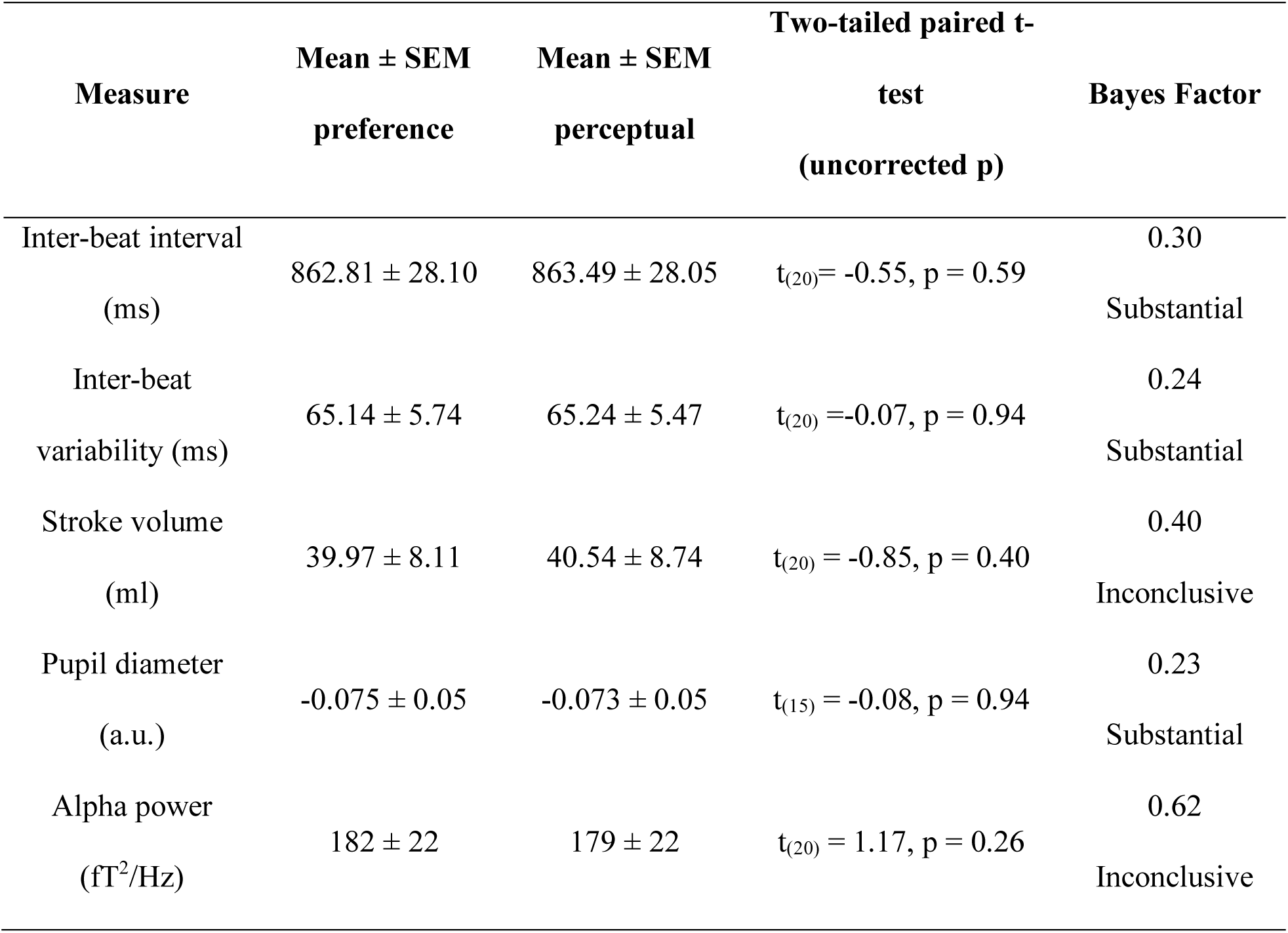
Cardiac parameters and arousal measures during instructions do not differ between preference-based and perceptual decisions.

### The subjective value of the chosen option is encoded in medial prefrontal cortices in preference-based decisions

We then identified when and where subjective value was encoded during preference-based choice. First, we modeled single trial response-locked neural activity at the sensor level using a GLM (GLM1a, see **Methods**), using as regressors the subjective values of the chosen (Chosen_SV_) and unchosen (Unchosen_SV_) options, as well as the response button used. Neural activity over frontal sensors encoded the subjective value of the chosen option in two neighboring time-windows (ß_ChosenSV_, first cluster: −580 to −370 ms before response, sum(t)=−7613, Monte Carlo p = 0.004. Second cluster: −336 to −197 ms before response, sum(t) = −4405, Monte Carlo p= 0.033; **Fig. 2D** and **Fig. 2E**). No cluster of neural activity significantly encoded the subjective value of the unchosen option. Motor preparation was encoded later in time in two posterior-parietal clusters of opposite polarities (ß_Button Press_, negative cluster: −287 to −28 ms before response, sum(t) = −10918, Monte Carlo p = 0.003; positive cluster: −373 to −196 ms before response, sum(t) = 5848, Monte Carlo p = 0.02).

To identify the cortical regions contributing to the encoding of subjective value at sensor-level, we used the same model (GLM1a) to predict source-reconstructed activity averaged in the time-window identified at sensor level (−580 to −197 ms before response). The subjective value of the chosen option was encoded as expected in medial prefrontal regions (right posterior vmPFC area 32 and 24, cluster peak at MNI coordinates: [7 40 0]; t-value = 4.52); right dorso-medial prefrontal cortex dmPFC area 8m [5 30 40]; t-value = 3.73), as well as in bilateral occipital poles ([6 −77 11], [−1 −85 16]; t-values, 4.17 and 3.85, respectively) and mid-posterior left insula ([−34 −27 17]; t-value = 4.48) (**Fig. 2F**).

### HER amplitude during instruction interacts with subjective value encoding in right vmPFC during choice on a trial-by-trial basis

We thus show that two different sub-regions of vmPFC were involved at different moments in a trial: during the instruction period, HERs were larger when participants prepared for preference-based vs. perceptual decisions in left and right anterior vmPFC, and during the choice period, subjective value was encoded in right posterior vmPFC. We then addressed our main question (**Fig. 1B**): does the amplitude of neural responses to heartbeats during the instruction period interact with the encoding of subjective value during choice in vmPFC?

We tested whether subjective value encoding in right posterior vmPFC was affected by HER amplitude measured in either left or right anterior vmPFC in a two-by-two ANOVA with HER amplitude (high or low, median split across trials) and hemisphere as factors. The ANOVA revealed a significant interaction between HER amplitude and hemisphere (**Fig. 3A**; F_(1,40)_=5.07, p=0.036; no main effect of HER amplitude F_(1,20)_=2.69, p=0.12, no main effect of hemisphere F_(1,20)_=0.19, p=0.67). This interaction corresponded to a significantly stronger subjective value encoding in trials where HERs in right vmPFC were larger during instructions (two-tailed paired t-test on ß_ChosenSV_ in large vs. small HER values in right vmPFC, t_(20)_ = 2.52, p = 0.02, Cohen’s d = 0.55).

**Figure 3.**
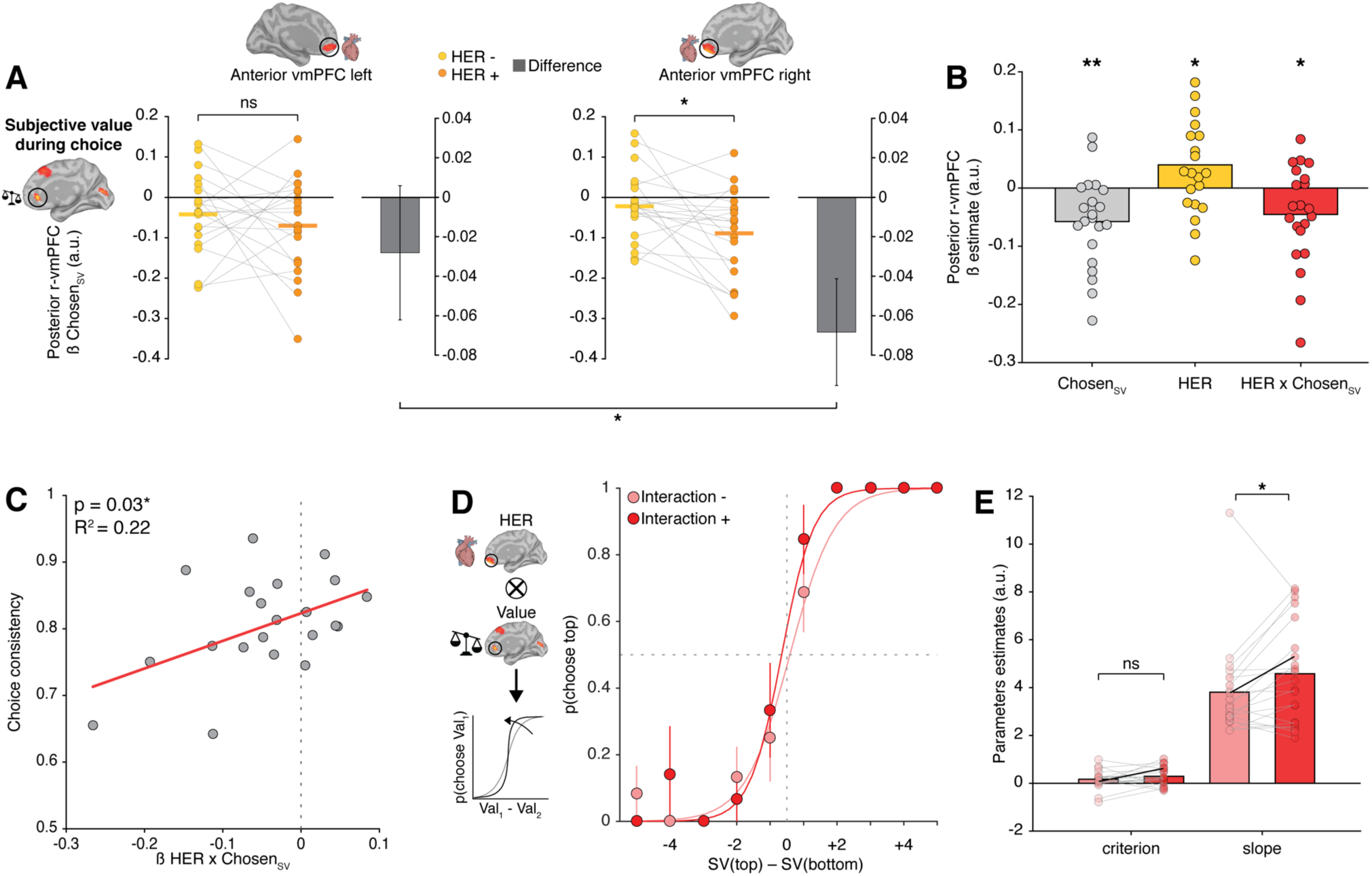
The interaction between HER and value encoding accounts for inter-individual variability in choice consistency and for intra-individual trial-by-trial fluctuations of choice precision. **A**, Parameter estimates for the encoding of the chosen option value in posterior r-vmPFC during choice, in trials where HERs during task preparation were small (yellow) or large (orange) in left anterior vmPFC (left), or in right anterior vmPFC (right). Each dot represents a participant, the horizontal line indicates the mean. The difference in value encoding between large and small HER (grey bars; error bars indicate SEM) was significant in right vmPFC (ANOVA on value encoding, significant interaction HER amplitude x hemisphere, F_(1,40)_=5.07 p=0.036); two tailed paired t-test comparing encoding strength for trials with large or small HERs in right vmPFC, t_(20)_=2.52 p=0.02) **B**, Activity in posterior r-vmPFC is by design explained by the chosen option subjective value, but it is also explained by HER amplitude in anterior r-vmPFC during instructions and by the interaction between HER and value encoding. **C**, Robust regression shows that the magnitude of the interaction between HER and value encoding positively predicts inter-individual variability in choice consistency **D**, Behavioral data (dots) and fitted psychometric function (lines) for one representative participant, in trials with a large or small interaction between HER and value encoding. Error bars represent SEM. **E**, Criterion and slope of the psychometric function in all participants, revealing a significantly steeper slope for trials with large interaction between HER and value-related vmPFC activity (p = 0.037). Decision criterion is unaffected. Black lines represent the parameters estimates of the participant displayed in D. *p<0.05, **p<0.01.

The influence of HER amplitude was specific to value encoding: the amplitude of visual responses evoked by option presentation was unrelated to HER amplitude (**Table 3**). HER amplitude is thus not a mere index of cortical responsiveness, interacting with any other brain response. HER amplitude in r-vmPFC did not vary with pupil diameter nor alpha power, neither during choice nor during value encoding (**Table 3**), indicating that HER fluctuations are not driven by an overall change in brain state. Last, to definitively rule out an influence of attention/arousal, we tested whether the strength of value encoding was modulated by fluctuations in alpha power or pupil diameter measured during instructions. We median-split trials based on either alpha power or pupil but found no difference in value encoding (alpha, paired t-test on ß_ChosenSV_ t_(20)_=0.19, p=0.85, BF=0.25; pupil, paired t-test t_(20)_=−0.23, p=0.82, BF=0.24).

**Table 3.** Arousal states and physiological parameters do not differ between low and high HER amplitude in preference trials.

| **Measure** | **Low HER**<br>**(Mean ± SEM)** | **High HER (Mean ±**<br>**SEM)** | **Two-tailed t-test**<br>**(uncorrected p)** | **Bayes**<br>**Factor** |
| --- | --- | --- | --- | --- |
| Pupil diameter (a.u.) during<br>option presentation, [-1 0] s | 0.27 ± 0.07 | 0.24 ± 0.07 | $t_{(15)} = 0.57$,<br>p = 0.58 | 0.29<br>Substantial |
| Pupil diameter (a.u.) during<br>value encoding<br>[-0.580 -0.197] s | 0.32 ± 0.08 | 0.29 ± 0.08 | $t_{(15)} = 0.57$,<br>p = 0.58 | 0.29<br>Substantial |
| Alpha power (fT<sup>2</sup>/Hz) max<br>15 channels during option<br>presentation, [-1 0] s | 1.24 x 10<sup>2</sup> ±<br>0.14 x 10<sup>2</sup> | 1.25 x 10<sup>2</sup> ± 0.13 x<br>10<sup>2</sup> | $t_{(20)} = -0.42$,<br>p = 0.68 | 0.27<br>Substantial |
| Alpha power (fT<sup>2</sup>/Hz) in the<br>value encoding cluster<br>[-0.580 -0.197] s | 0.56 x 10<sup>2</sup> ±<br>0.03 x 10<sup>2</sup> | 0.56 x 10<sup>2</sup> ± 0.03 x<br>10<sup>2</sup> | $t_{(20)} = -0.71$,<br>p = 0.48 | 0.35<br>Inconclusive |
| RMS of visual response to<br>options (fT) [0-250] ms | 62.8 ± 2.43 | 62.5 ± 2.40 | $t_{(20)} = 0.28$<br>p = 0.78 | 0.25<br>Substantial |

The interaction between HER amplitude and subjective value encoding in right vmPFC was further tested using a full parametric approach. Here (GLM2), we predicted the activity of posterior r-vmPFC during choice from the subjective value of the chosen option, the HER amplitude in anterior r-vmPFC during instruction period and the interaction between these two terms (**Fig. 3B**). Since the posterior vmPFC region of interest was defined based on its encoding of the chosen value, the parameter estimate for chosen value was, as expected, large (ß_ChosenSV_ = −0.06 ± 0.02, two-tailed t-test against 0, t_(20)_ = −3.37, p = 0.003). Activity in posterior vmPFC was also predicted by the amplitude of HERs occurring about 1.5 s earlier, during the instruction period, independently from the chosen value (ß_HER_ = 0.04 ± 0.02, two-tailed t-test against 0, t_(20)_= 2.13, p = 0.046), and importantly by the interaction between HERs and chosen value (ß_HER* ChosenSV_ = −0.05 ± 0.02, two-tailed t-test against 0, t_(20)_= −2.41, p = 0.025). Both the median-split analysis and the parametric model thus reveal a significant interaction between the amplitude of HERs during instruction period and the neural encoding of subjective value during choice.

We then verified that the effect on the neural encoding of subjective value was specific to HER amplitude, and not due to an overall baseline shift in anterior r-vmPFC during the instruction period. We ran an alternative model (GLM3) in which the activity in posterior r-vmPFC was predicted from the subjective value of the chosen option, the activity in anterior r-vmPFC averaged during the whole instruction period, i.e. activity not time-locked to heartbeats, and the interaction between the two terms. This analysis revealed that while the subjective value of the chosen option still significantly predicted the activity of posterior r-vmPFC (ß_ChosenSV_ = −0.05 ± 0.02, two-tailed t-test against 0, t_(20)_ = −3.27, p = 0.004), the other two terms did not (activity in anterior r-vmPFC averaged during instruction period: ß_BL vmPFC_ = 0.006 ± 0.03, two-tailed t-test against 0, t_(20)_ = 0.22, p = 0.83, BF = 0.25; interaction: ß_BL_ vmPFC* ChosenSV −0.03 ± 0.02, two-tailed t-test against 0, t_(20)_ = −1.55, p = 0.14, BF = 1.14). The encoding of subjective value is thus specifically modulated by HER amplitude in anterior r-vmPFC and not by an overall baseline shift unrelated to heartbeats in the same region.

The functional coupling between HERs and subjective value encoding was also region specific: HER amplitude in anterior r-vmPFC was unrelated to the strength of value encoding in any other value-related regions (two-tailed paired t-test on ß_ChosenSV_ in large vs. small HER values in right dmPFC, t_(20)_ = −0.89, p = 0.38, BF = 0.43; right occipital pole, t_(20)_ = −0.86, p = 0.40, BF = 0.41; left occipital pole, t_(20)_ = −1.60, p = 0.13, BF = 0.81; left posterior insula, t_(20)_= 1.00, p = 0.33, BF = 0.49). Conversely, HERs outside anterior r-vmPFC did not significantly interact with value encoding in posterior r-vmPFC. Splitting trials based on the amplitude in the two other cortical regions showing differential heartbeats-evoked responses (**Fig. 2C**) showed no significant modulation of value encoding in right posterior vmPFC (post-central complex: two-tailed paired t-test on ß_ChosenSV_, t_(20)_= −1.41, p = 0.17, BF = 0.90; right supramarginal gyrus: two-tailed paired t-test, t_(20)_= −1.96, p = 0.06, BF = 2.41).

### The interaction between HER and value encoding predicts choice consistency

To what extent does the interaction between HER and value encoding in vmPFC predict behavior? We first tested whether the interaction between HER and value encoding relates to inter-individual differences in choice consistency, i.e. whether participants selected the movie to which they had attributed the greatest likeability rating the day before. Given the overall high consistency in preference-based decisions, which may reduce our ability to detect significant relationships, we computed mean choice consistency using the top-50% most difficult trials (i.e. trials above median difficulty in each participant). We regressed the model parameter of the interaction between HER and value encoding (ß_HER*ChosenSV_ obtained from GLM2) against mean choice consistency across participants. The larger the interaction between HER and value encoding, the more consistent were participants in their choices (ß_robust_ = 0.41, robust regression R^2^ = 0.22, t_(19)_ = 2.29, p = 0.03; **Fig. 3C**). In other words, 22% of inter-individual difference in behavioral consistency is explained by the magnitude of the interaction between HER and value encoding.

The correlation between neural activity and behavior was specific to the interaction parameter: inter-individual differences in choice consistency could not be predicted from the model parameter estimate of HER (ß_HER_ from GLM2; ß_robust_ = 0.02, R^2^ = 4*10^−4^, t_(19)_ = 0.09, p = 0.93, BF = 0.39), nor from the parameter estimate of value (ß_ChosenSV_ from GLM2; ß_robust_ = −0.19, R^2^ = 0.04, t_(19)_ = −0.88, p = 0.39, BF = 0.52). The interaction between HER and subjective value encoding did not co-vary significantly neither with the personality traits, assessed through self-reported questionnaires (robust regressions on BDI scores, t_(19)_ = −0.17, p = 0.87, BF = 0.40; STAI scores, t_(19)_ = 0.90, p = 0.38, BF = 0.52; OCI scores, t_(19)_ = −1.23, p = 0.22, BF = 0.70; PDI scores, t_(19)_ = −0.32, p = 0.75, BF = 0.60) nor interoceptive ability, assessed with the heartbeat counting task (t_(19)_ = −0.73,p = 0.48, BF = 0.48).

So far, results are based on parameter estimates computed across trials for a given participant. To assess how within-participant trial-by-trial fluctuations in behavior relates to the interaction between HERs and subjective value encoding, we computed the z-scored product of the HER amplitude in anterior r-vmPFC during the instruction period and the value-related activity in posterior r-vmPFC during choice. We then median-split the trials according to this product and modeled participants’ choices separately for trials with a small vs. large interaction (**Fig. 3D**). When the interaction was large, psychometric curves featured a steeper slope, corresponding to an increased choice precision (two-tailed paired t-test, t_(20)_= −2.24, p = 0.037, Cohen’s d = −0.49; after removal of the unique outlier with a slope estimate exceeding 3 SD from population mean, t_(19)_ = −3.30, p = 0.003, Cohen’s d= −0.74; **Fig. 3E**), while decision criterion was not affected (two-tailed paired t-test, t_(20)_= −1.20, p = 0.25, BF = 0.64; after outlier removal t_(19)_ =−0.96, p = 0.35, BF = 0.46; **Fig. 3E**).

To control for the specificity of the interaction, we estimated the psychometric function on trials median-split on HER amplitude alone but found no difference in choice precision (two-tailed paired t-test, t_(20)_= 0.41, p = 0.69, BF = 0.27) nor in criterion (two-tailed paired t-test, t_(20)_= 0.52, p = 0.61, BF = 0.29). Similarly, median-splitting trials on value-related posterior r-vmPFC activity alone revealed no difference in the psychometric curve (two-tailed paired t-test, slope, t_(20)_= 0.21, p = 0.84, BF = 0.25; criterion: t_(20)_= −0.57, p = 0.58, BF = 0.30). Therefore, our results indicate that trial-by-trial choice precision is specifically related to the interaction between HERs in anterior r-vmPFC and value-related activity in posterior r-vmPFC.

Altogether, these results indicate that the interaction between HER amplitude and subjective value encoding accounts both for within-subject inter-trial variability, and for inter-individual differences in preference-based choice consistency.

### HER effects are specific to preference-based choices

Finally, we tested whether the effect of HER was specific to subjective value or whether it is a more general mechanism interacting with all types of decision-relevant evidence. To this aim, we analyzed perceptual discrimination trials using the same approach as for preference-based trials. First, we modeled the single trial response-locked MEG sensor-level data using a GLM (GLM1b) with the parameters accounting for choice in the perceptual task (i.e. contrast of the chosen option – Chosen_Ctrs_ – and the contrast of the unchosen option – Unchosen_Ctrs_), as well as response button. The non-parametric clustering procedure revealed the presence of a fronto-central cluster encoding the contrast of the chosen option (ß_ChosenCtrs_, −257 to −25 ms before response, sum(t)= 8121, Monte Carlo p = 0.005; **Fig. 4A** and **B)**. We also found two clusters of opposite polarities encoding the contrast of the unchosen option (ß_UnchosenCtrs,_ positive cluster, −250 to −79 ms before response, sum(t) = 6182, Monte Carlo p = 0.008; negative cluster, −211 to −88 ms before response, sum(t) = −4127, Monte Carlo p = 0.04) and two clusters encoding motor preparation (ß_Button Press,_ positive cluster, −193 to 0 ms before response, sum(t) = 11103, Monte Carlo p = 0.0004; negative cluster, −222 to 0 ms before response, sum(t) = −10395, Monte Carlo p = 0.0004). The same model (GLM1b) applied to source-reconstructed activity averaged in the time-window identified at sensor level (−257 to −25 ms before response) revealed four cortical areas encoding the contrast of the chosen option (**Fig. 4C**): left midcingulate area (peak at MNI coordinates [−11 −27 43], t-value = 5.14), left superior frontal gyrus ([−15 7 71], t-value = 4.70) and bilateral inferior parietal lobule (right IPL, [−47 −55 53], t-value = 4.25; left IPL, [44 −42 49], t-value = 5.51). Finally, we median-split perceptual trials according to the amplitude of HERs in anterior r-vmPFC. The encoding strength of the contrast of the chosen option did not interact with heartbeat-evoked responses amplitude in any of the contrast-encoding regions (all p ≥ 0.26, BF ≤ 0.62; **Table 4**). The results thus indicate that HER amplitude in r-vmPFC is specifically linked to the cortical encoding of subjective value.

**Table 4.**
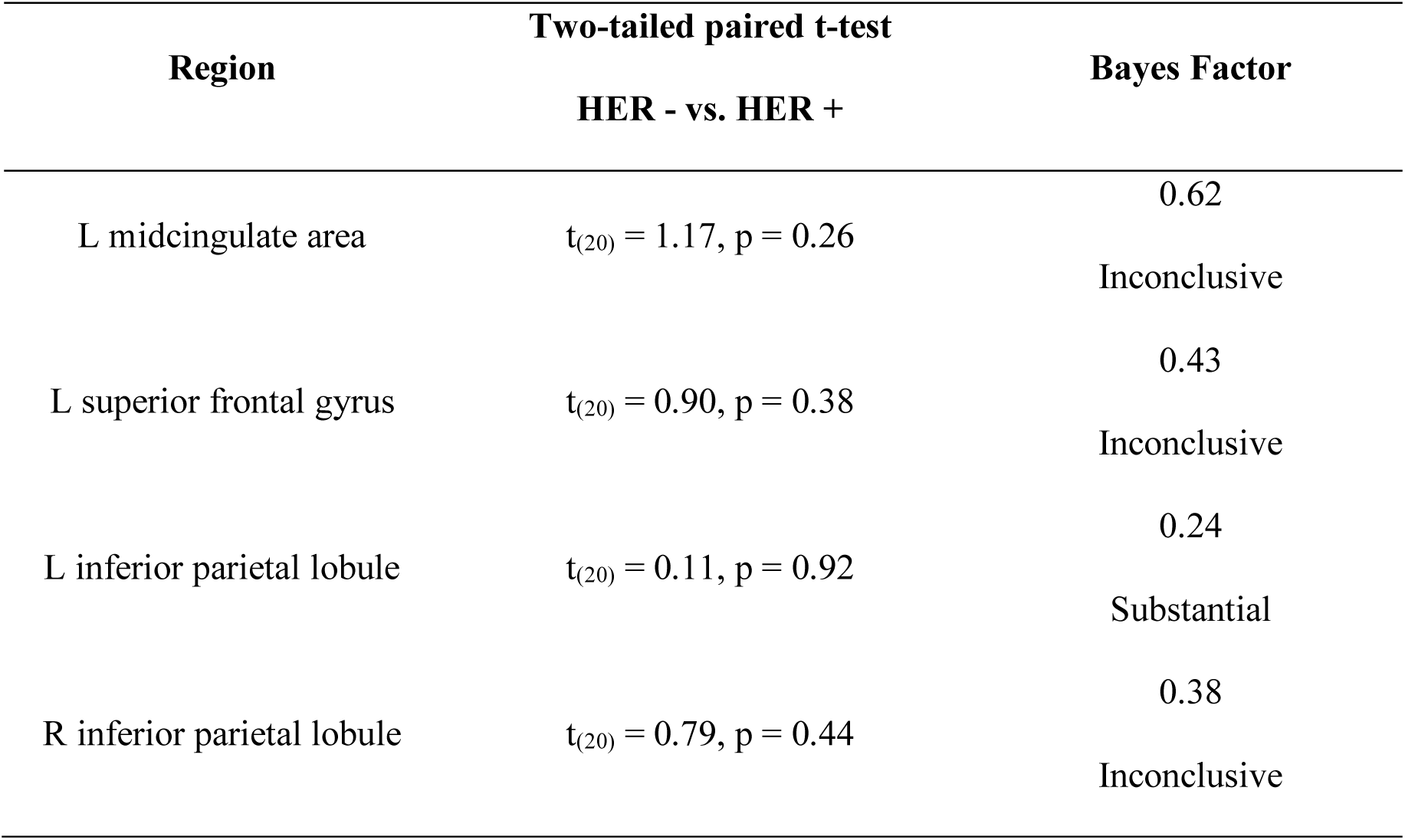
Encoding strength for contrast during perceptual decisions does not depend on HER amplitude in anterior r-vmPFC.

**Figure 4.**
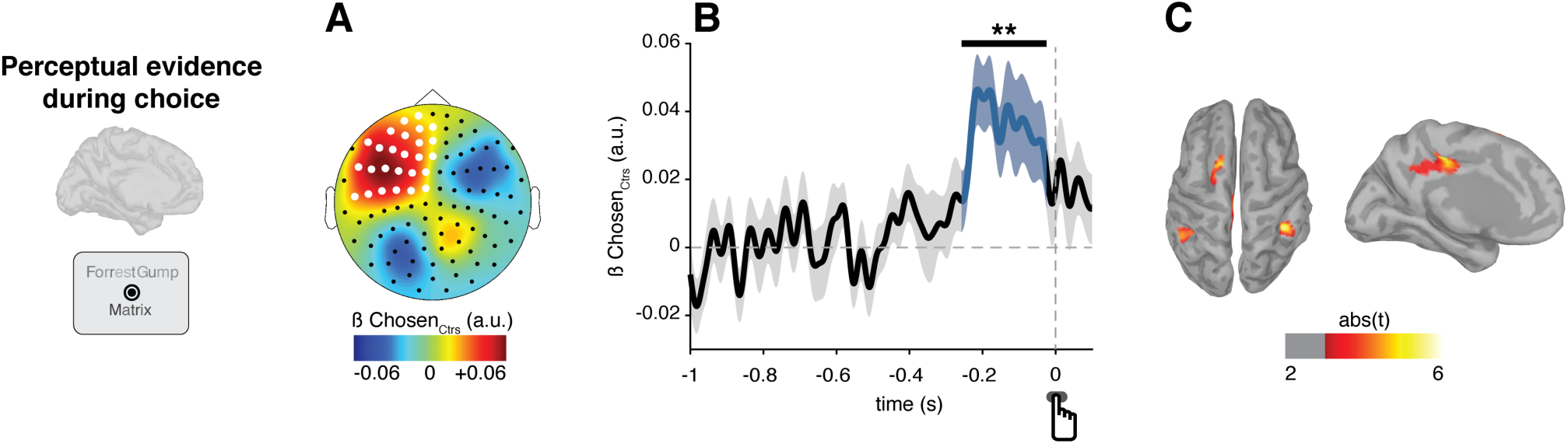
Neural encoding of perceptual evidence. **A**, Topography of the significant encoding of the chosen option contrast (−257 to −25 ms before motor response) during choice in objective visual discrimination trials. **B**, Time-course (± SEM) of the GLM parameter estimate for the chosen option contrast in the cluster highlighted in white in A. The black bar indicates the significant time-window, as established by non-parametric clustering procedure. **C**, Brain regions mostly contributing to the encoding of the contrast of the chosen option (at least 20 contiguous vertices with uncorrected p < 0.005). **p<0.01.

## Discussion

We show that preparing for subjective preference-based decisions led to larger responses to heartbeats in vmPFC, as expected from previous studies relating self and HERs (Park et al., 2014; Babo-Rebelo et al., 2016a, 2016b), and in post-central gyrus, a region known to respond to heartbeats (Kern et al., 2013; Azzalini et al., 2019; Al et al., 2020). We further reveal that HERs before option presentation interact specifically with subjective value encoding during choice in vmPFC. The interaction between HERs and value encoding predicted preference-based choices: it was associated to more precise decisions at the single trial level, and predicted inter-individual variability in choice consistency over time. No interaction between HER and the encoding of perceptual evidence could be found in the control objective task. Neither the HER difference between the two tasks, nor the interaction between HER and value in the subjective preference task, could be trivially explained by changes in cardiac parameters (heart rate, heart rate variability, ECG, stroke volume) nor by changes in arousal state (pupil diameter, alpha power). Altogether, our results reveal how the self-reflection intrinsic to preference-based decisions involves the neural read-out of a physiological variable and its integration in the subjective valuation process.

In line with previous studies relating the self with HERs in vmPFC (Park et al., 2014; Babo-Rebelo et al., 2016a, 2016b), we find that HERs are more pronounced when preparing for the subjective preference task than when preparing for the objective discrimination task. A number of self-related processes might take place specifically when preparing for the subjective preference-based task, such as turning attention inward or pre-activating autobiographical memory circuits. Note that HERs cannot be influenced by processes triggered by the movie titles themselves, such as retrieving movie-specific information from memory, since options are not yet available in the instruction period where HERs are measured. HERs were not associated with any change in the neural encoding of objective perceptual evidence. This result might seem at odds with previously reported effects of HERs on sensory detection at threshold (Park et al., 2014; Al et al., 2020). However, as opposed to the perceptual discrimination task used here, sensory detection at threshold is intrinsically subjective, since participants are asked to introspect and report their fluctuating subjective experience in response to physically and objectively constant stimuli (Ress and Heeger, 2003; Campana and Tallon-Baudry, 2013). Last, there was no difference in cardiac parameters between the two tasks. Our results thus differ from previous observations in risky decisions, where changes in peripheral bodily signals predicted behavioral performance (Bechara et al., 1997). More generally, this indicates that HER fluctuations relevant for valuation and behavior correspond to changes in the quality of the neural monitoring in cardiac inputs, rather than to changes in cardiac activity.

We successfully retrieved with MEG the cortical valuation network described in the fMRI literature (Levy and Glimcher, 2012; Bartra et al., 2013; Clithero and Rangel, 2014), including dmPFC and vmPFC, during the choice period. These findings are interesting *per se*, as data on the temporal course of value-based choices in the human prefrontal cortex remain scarce (Hunt et al., 2012, 2015; Polanía et al., 2014; Lopez-Persem et al., 2020). Here, we find that chosen value is robustly encoded in the valuation network from 600 ms to 200 ms before motor response, with a temporal (but not spatial) overlap with motor preparation. Note that we did not find a robust encoding of the unchosen value, in line with electrophysiological recordings in the monkey orbitofrontal cortex where the encoding of the chosen value dominates (Padoa-Schioppa and Assad, 2006; Strait et al., 2014; Hunt et al., 2015; Rich and Wallis, 2016). Chosen value encoding interacted with HERs specifically in vmPFC, a region shown by separate streams of studies to play a role in valuation (Fellows, 2006; Delgado et al., 2016; Vaidya et al., 2017) and self-related cognition (Qin and Northoff, 2011; Andrews-Hanna et al., 2014), and long suspected to play an integrative role in decision making (Vaidya and Fellows, 2020). In the objective discrimination task, relevant perceptual evidence was encoded, among other regions, in posterior parietal cortex, consistent with monkey electrophysiology literature (Shadlen and Newsome, 2001; Heekeren et al., 2008).

Because fluctuations in HERs occurred during task preparation, before option presentation, their influence on value encoding might generally pertain to interactions between ongoing activity (during task preparation) and stimulus-evoked activity (in response to option presentation). HERs constitute a very specific subset of such interactions: HERs interacted only with value, but not with visual evoked responses for instance, and only in vmPFC in the subjective task. Conversely, no interaction between overall ongoing activity in vmPFC and value encoding could be found. Besides, HERs modulated value encoding in a multiplicative manner. Our results thus differ from a previously reported vmPFC baseline-shift additive effect in pleasantness ratings (Abitbol et al., 2015; Lopez-Persem et al., 2020). Altogether, our results indicate that part of the unspecified ‘neural noise’ driving fluctuations in choice consistency (Padoa-Schioppa, 2013; Kurtz-David et al., 2019; Webb et al., 2019) comes from the interaction between interoceptive self-related processes, indexed by neural monitoring of cardiac signals, and the neural encoding of subjective value. A more detailed mechanistic account of how responses to heartbeat during task preparation influence the subjective valuation process taking place about 1.5 second later remains to be established. Still, our results pave the way for the quantification of self-related processes in decision making, an aspect mostly absent from computational models of decision-making despite its relevance to understand maladaptive decisions in psychiatric disorders (Paulus, 2007; Moeller and Goldstein, 2014; Sui and Gu, 2017).

Decisions on primary goods such as food integrate information about internal bodily states to select options that preserve the integrity of the organism that needs to be fed and protected – the simplest notion of self. We show here that the subjective valuation of cultural goods, that relies on the same cortical valuation network as employed for primary goods (Chib et al., 2009; Lebreton et al., 2009; Levy and Glimcher, 2011; McNamee et al., 2013; Sescousse et al., 2013), has inherited a functional link with the central monitoring of physiological variables. Even when choosing between cultural goods that do not fulfill any immediate basic need, the neural monitoring of heartbeats supports self-reflection underlying evaluation, contributing to the precision of subjective decisions and fostering the stable expression of long-lasting preferences that define, at least in part, our identity.

## Acknowledgements

This work was supported by funding from the European Research Council (ERC) under the European Union’s Horizon 2020 research and innovation program (grant agreement No 670325, Advanced grant BRAVIUS) and a senior fellowship of the Canadian Institute For Advanced Research (CIFAR) program in Brain, Mind and Consciousness to CTB; by a doctoral fellowship from the Ecole des Neurosciences de Paris Ile de France to DA; and by ANR-17-EURE-0017.

The authors thank Clémence Alméras, Maximilien Chaumon and Christophe Gitton for assistance in data acquisition.

## References

Abitbol R, Lebreton M, Hollard G, Richmond BJ, Bouret S, Pessiglione M (2015) Neural Mechanisms Underlying Contextual Dependency of Subjective Values: Converging Evidence from Monkeys and Humans. J Neurosci 35:2308–2320.

Al E, Iliopoulos F, Forschack N, Nierhaus T, Grund M, Motyka P, Gaebler M, Nikulin V V., Villringer A (2020) Heart-brain interactions shape somatosensory perception and evoked potentials. Proc Natl Acad Sci U S A 117:10575–10584.

Andrews-Hanna JR, Smallwood J, Spreng RN (2014) The default network and self-generated thought: Component processes, dynamic control, and clinical relevance. Ann N Y Acad Sci 1316:29–52.

Azzalini D, Rebollo I, Tallon-Baudry C (2019) Visceral Signals Shape Brain Dynamics and Cognition. Trends Cogn Sci 23:488–509.

Babo-Rebelo M, Richter CG, Tallon-Baudry C (2016a) Neural Responses to Heartbeats in the Default Network Encode the Self in Spontaneous Thoughts. J Neurosci 36:7829–7840.

Babo-Rebelo M, Wolpert N, Adam C, Hasboun D, Tallon-Baudry C (2016b) Is the cardiac monitoring function related to the self in both the default network and right anterior insula? Philos Trans R Soc B Biol Sci 371:20160004.

Bartra O, McGuire JT, Kable JW (2013) The valuation system: A coordinate-based meta-analysis of BOLD fMRI experiments examining neural correlates of subjective value. Neuroimage 76:412–427.

Bechara A, Damasio H, Tranel D, Damasio AR (1997) Deciding Advantageously Before Knowing the Advantageous Strategy. Science (80-) 275:1293–1295.

Beck A, Ward CH, Mendelson M, Mock J, Erbaught J (1961) An Inventory for Measuring Depression. Arch Gen Psychiatry 4:561–571.

Berntson GG, Quigley KS, Lozano D (2007) Cardiovascular Psychophysiology. In: Handbook of Psychophysiology, 3rd ed. (Cacioppo JT, Tassinary LG, Berntson G, eds), pp 182–210. Cambridge University Press.

Blanke O, Metzinger T (2009) Full-body illusions and minimal phenomenal selfhood. Trends Cogn Sci 13:7–13.

Campana F, Tallon-Baudry C (2013) Anchoring visual subjective experience in a neural model: The coarse vividness hypothesis. Neuropsychologia 51:1050–1060.

Chib VS, Rangel A, Shimojo S, O’Doherty JP (2009) Evidence for a Common Representation of Decision Values for Dissimilar Goods in Human Ventromedial Prefrontal Cortex. J Neurosci 29:12315–12320.

Clithero JA, Rangel A (2014) Informatic parcellation of the network involved in the computation of subjective value. Soc Cogn Affect Neurosci 9:1289–1302.

Craig AD (2002) How do you feel? Interoception: the sense of the physiological condition of the body. Nat Rev Neurosci 3:655–666.

Damasio AR (2010) Self Comes to Mind: Constructing the Conscious Brain. Pantheon Books.

Delgado MR, Beer JS, Fellows LK, Huettel SA, Platt ML, Quirk GJ, Schiller D (2016) Viewpoints: Dialogues on the functional role of the ventromedial prefrontal cortex. Nat Neurosci 19:1545–1552.

Delorme A, Makeig S (2004) EEGLAB: an open source toolbox for analysis of single-trial EEG dynamics including independent component analysis. J Neurosci Methods 134:9–21.

Desmedt O, Luminet O, Corneille O (2018) The heartbeat counting task largely involves non-interoceptive processes: Evidence from both the original and an adapted counting task. Biol Psychol 138:185–188.

Dirlich G, Vogl L, Plaschke M, Strian F (1997) Cardiac field effects on the EEG. Electroencephalogr Clin Neurophysiol 102:307–315.

Fellows LK (2006) Deciding how to decide: Ventromedial frontal lobe damage affects information acquisition in multi-attribute decision making. Brain 129:944–952.

Fischl B, Van Der Kouwe A, Destrieux C, Halgren E, Ségonne F, Salat DH, Busa E, Seidman LJ, Goldstein J, Kennedy D, Caviness V, Makris N, Rosen B, Dale AM (2004) Automatically Parcellating the Human Cerebral Cortex. Cereb Cortex 14:11–22.

Foa EB, Huppert JD, Leiberg S, Langner R, Kichic R, Hajcak G, Salkovskis PM (2002) The Obsessive-Compulsive Inventory: Development and validation of a short version. Psychol Assess 14:485–496.

Gray MA, Taggart P, Sutton PM, Groves D, Holdright DR, Bradbury D, Brull D, Critchley HD (2007) A cortical potential reflecting cardiac function. PNAS 104:6818–6823.

Grueschow M, Polania R, Hare TA, Ruff CC (2015) Automatic versus Choice-Dependent Value Representations in the Human Brain. Neuron 85:874–885.

Heekeren HR, Marrett S, Ungerleider LG (2008) The neural systems that mediate human perceptual decision making. Nat Rev Neurosci 9:467–479.

Hunt LT, Behrens TEJ, Hosokawa T, Wallis JD, Kennerley SW (2015) Capturing the temporal evolution of choice across prefrontal cortex. Elife 4:1–25.

Hunt LT, Kolling N, Soltani A, Woolrich MW, Rushworth MFS, Behrens TEJ (2012) Mechanisms underlying cortical activity during value-guided choice. Nat Neurosci 15:470–476.

Juechems K, Summerfield C (2019) Where does value come from? Trends Cogn Sci:836–850.

Kass RE, Raftery AE (1995) Bayes factors. J Am Stat Assoc 90:773–795.

Keramati M, Gutkin B (2014) Homeostatic reinforcement learning for integrating reward collection and physiological stability. Elife 3:1–26.

Kern M, Aertsen A, Schulze-Bonhage A, Ball T (2013) Heart cycle-related effects on event-related potentials, spectral power changes, and connectivity patterns in the human ECoG. Neuroimage 81:178–190.

Kubicek WG, Patterson RP, Witsoe DA (1970) Impedance Cardiography As a Noninvasive Method of Monitoring Cardiac Function and Other Parameters of the Cardiovascular System. Ann N Y Acad Sci 170:724–732.

Kurtz-David V, Persitz D, Webb R, Levy DJ (2019) The neural computation of inconsistent choice behavior. Nat Commun 10:1583.

Lebreton M, Jorge S, Michel V, Thirion B, Pessiglione M (2009) An Automatic Valuation System in the Human Brain: Evidence from Functional Neuroimaging. Neuron 64:431–439.

Levy DJ, Glimcher PW (2011) Comparing Apples and Oranges: Using Reward-Specific and Reward-General Subjective Value Representation in the Brain. J Neurosci 31:14693–14707.

Levy DJ, Glimcher PW (2012) The root of all value: A neural common currency for choice. Curr Opin Neurobiol 22:1027–1038.

Liang F, Paulo R, Molina G, Clyde MA, Berger JO (2008) Mixtures of g priors for Bayesian variable selection. J Am Stat Assoc 103:410–423.

Lopez-Persem A, Bastin J, Petton M, Abitbol R, Lehongre K, Adam C, Navarro V, Rheims S, Kahane P, Domenech P, Pessiglione M (2020) Four core properties of the human brain valuation system demonstrated in intracranial signals. Nat Neurosci 23:664–675.

Maris E, Oostenveld R (2007) Nonparametric statistical testing of EEG- and MEG-data. J Neurosci Methods 164:177–190.

McNamee D, Rangel A, O’Doherty JP (2013) Category-dependent and category-independent goal-value codes in human ventromedial prefrontal cortex. Nat Neurosci 16:479–485.

Moeller SJ, Goldstein RZ (2014) Impaired self-awareness in human addiction: deficient attribution of personal relevance. Trends Cogn Sci 18:635–641.

Montoya P, Schandry R, Müller A (1993) Heartbeat evoked potentials (HEP): topography and influence of cardiac awareness and focus of attention. Electroencephalogr Clin Neurophysiol Evoked Potentials 88:163–172.

Neubert F-X, Mars RB, Sallet J, Rushworth MFS (2015) Connectivity reveals relationship of brain areas for reward-guided learning and decision making in human and monkey frontal cortex. Proc Natl Acad Sci U S A 112:E2695–704.

Oostenveld R, Fries P, Maris E, Schoffelen JM (2011) FieldTrip: Open source software for advanced analysis of MEG, EEG, and invasive electrophysiological data. Comput Intell Neurosci 2011:1–9.

Padoa-Schioppa C (2013) Neuronal origins of choice variability in economic decisions. Neuron 80:1322–1336.

Padoa-Schioppa C, Assad JA (2006) Neurons in the orbitofrontal cortex encode economic value. Nature 441:223–226.

Park H-D, Correia S, Ducorps A, Tallon-Baudry C (2014) Spontaneous fluctuations in neural responses to heartbeats predict visual detection. Nat Neurosci 17:612–618.

Park H-D, Tallon-Baudry C (2014) The neural subjective frame: from bodily signals to perceptual consciousness. Philos Trans R Soc Lond B Biol Sci 369:20130208.

Paulus MP (2007) Decision-Making Dysfunctions in Psychiatry—Altered Homeostatic Processing? Science (80-) 318:602–606.

Peters E, Joseph S, Day S, Garety P (2004) Measuring Delusional Ideation: The 21-Item Peters et aL Delusions Inventory (PDI). Schizophr Bull 30:1005–1022.

Polanía R, Krajbich I, Grueschow M, Ruff CC (2014) Neural Oscillations and Synchronization Differentially Support Evidence Accumulation in Perceptual and Value-Based Decision Making. Neuron 82:709–720.

Qin P, Northoff G (2011) How is our self related to midline regions and the default-mode network? Neuroimage 57:1221–1233.

Ress D, Heeger DJ (2003) Neuronal correlates of perception in early visual cortex. Nat Neurosci 6:414–420.

Rich EL, Wallis JD (2016) Decoding subjective decisions from orbitofrontal cortex. Nat Neurosci 19.

Ring C, Brener J, Knapp K, Mailloux J (2015) Effects of heartbeat feedback on beliefs about heart rate and heartbeat counting: A cautionary tale about interoceptive awareness. Biol Psychol 104:193–198.

Schandry R (1981) Heartbeat perception and emotional experience. Psychophysiology 18:483–488.

Sescousse G, Caldú X, Segura B, Dreher JC (2013) Processing of primary and secondary rewards: A quantitative meta-analysis and review of human functional neuroimaging studies. Neurosci Biobehav Rev 37:681–696.

Shadlen MN, Newsome WT (2001) Neural basis of a perceptual decision in the parietal cortex (area LIP) of the rhesus monkey. J Neurophysiol 86:1916–1936.

Sherwood A, Allen M, Fahrenberg J, Kelsey RM, Lovallo WR, van Doornen LJP (1990) Methodological Guidelines for Impedance Cardiography. Psychophysiology 27:1–23.

Spielberger CD, Gorsuch RL, Lushene R, Vagg PR, Jacobs GA (1983) Manual for the State-Trait Anxiety Inventory. In: Consulting Psychologists. Palo Alto, California.

Strait CE, Blanchard TC, Hayden BY (2014) Reward value comparison via mutual inhibition in ventromedial prefrontal cortex. Neuron 82:1357–1366.

Sui J, Gu X (2017) Self as Object: Emerging Trends in Self Research. Trends Neurosci 40:643–653.

Tadel F, Baillet S, Mosher JC, Pantazis D, Leahy RM (2011) Brainstorm: A user-friendly application for MEG/EEG analysis. Comput Intell Neurosci 2011:879716.

Tzourio-Mazoyer N, Landeau B, Papathanassiou D, Crivello F, Etard O, Delcroix N, Mazoyer B, Joliot M (2002) Automated anatomical labeling of activations in SPM using a macroscopic anatomical parcellation of the MNI MRI single-subject brain. Neuroimage 15:273–289.

Vaidya AR, Fellows LK (2020) Under construction: ventral and lateral frontal lobe contributions to value-based decision-making and learning. F1000Research 9:1–8.

Vaidya AR, Sefranek M, Fellows LK (2017) Ventromedial Frontal Lobe Damage Alters how Specific Attributes are Weighed in Subjective Valuation. Cereb Cortex 28:3857–3867.

Vinck M, van Wingerden M, Womelsdorf T, Fries P, Pennartz CMA (2010) The pairwise phase consistency: A bias-free measure of rhythmic neuronal synchronization. Neuroimage 51:112–122.

Webb R, Levy I, Lazzaro SC, Rutledge RB, Glimcher PW (2019) Neural random utility: Relating cardinal neural observables to stochastic choice behavior. J Neurosci Psychol Econ 12:45–72.

Zamariola G, Maurage P, Luminet O, Corneille O (2018) Interoceptive accuracy scores from the heartbeat counting task are problematic: Evidence from simple bivariate correlations. Biol Psychol 137:12–17.

